# NR2F1 is a barrier to dissemination of early-evolved mammary cancer cells

**DOI:** 10.1101/2021.01.29.428822

**Authors:** Carolina Rodriguez-Tirado, Nupura Kale, Maria Jose Carlini, Nitisha Shrivastava, Bassem Khalil, Jose Javier Bravo-Cordero, Melissa Alexander, Jiayi Ji, Maria Soledad Sosa

## Abstract

Cancer cells disseminate during very early and sometimes asymptomatic stages of tumor progression. Granted that biological barriers to tumorigenesis exist, there must also be limiting steps to early dissemination, all of which remain largely unknown. We report that the orphan nuclear receptor NR2F1/COUP-TF1 serves as a barrier to early dissemination. High-resolution intravital imaging revealed that loss of function of NR2F1 in HER2+ early cancer cells increased *in vivo* dissemination without accelerating mammary tumor formation. NR2F1 expression was positively regulated by the tumor suppressive MMK3/6-p38-MAPK pathway and downregulated by HER2 and Wnt4 oncogenic signaling. NR2F1 downregulation by HER2 in early cancer cells led to decreased E-cadherin expression and β-catenin membrane localization, disorganized laminin 5 deposition, and increased expression of CK14, TWIST1, ZEB1 and PRRX1. Our findings reveal the existence of an inhibitory mechanism of dissemination regulated by NR2F1 downstream of HER2 signaling.

## Introduction

Accumulating evidence in experimental models and human specimens across different cancers supports that early in cancer evolution, cancer cells are able to disseminate to different anatomical sites^1–11^. For instance, patients with breast ductal carcinoma in situ (DCIS) lesions defined by pathologists as non-invasive carry early disseminated cancer cells (eDCCs) in target organs^5^. Interestingly, a small, but not less significant, percentage of women with DCIS who never progressed to an in-breast invasive disease died from a distant recurrence^8,12^ suggesting that eDCCs can fuel the formation of metastatic events. However, the mechanisms responsible for early cancer cells (ECCs) to acquire motile and invasive features allowing dissemination and target organ colonization are not clearly understood.

We previously showed in the MMTV-HER2 mammary cancer mouse model, that the early activation of the HER2 oncogene in mammary epithelial cells induced a deregulated branching morphogenesis program by turning off p38α/β signaling and promoting a Wnt-dependent partial EMT program^13^. This allowed ECCs to acquire an invasive phenotype and disseminate to secondary organs. We previously showed that epithelial cancer cells (e.g. head and neck squamous cell carcinoma, HNSCC) commonly downregulate the expression of a member of the steroid hormone receptor family NR2F1 (COUP-TF1/Ear3) and that upregulation of p38α/β signaling could restore NR2F1 expression^14^. NR2F1 levels were downregulated in early lesions and primary tumors from the MMTV-Myc and MMTV-HER2 mouse models when compared to wild type siblings^14^. Then, the arising question is whether the loss of NR2F1 in early lesions is linked to tumor suppression, early dissemination or both.

Here we show that in murine and human early mammary cancer lesions, HER2 signaling reduces NR2F1 expression and this requires reduced MKK3/6-p38-MAPK activity. We also demonstrate that NR2F1 downregulation in HER2+ ECCs resulted in disorganized laminin V, reduced E-cadherin expression and total membrane β-catenin localization as well as gain of basal CK14+ phenotype in invading cells. Importantly, NR2F1 depletion in ECCs also facilitated the acquisition of partial EMT program components downstream of HER2, which included the upregulation of TWIST1, *Zeb1* and PRRX1 without any changes in VIMENTIN, SNAIL and *Axin2*. We also showed using high-resolution intravital two-photon microscopy that loss of NR2F1 in MMTV-HER2 ECCs favored *in vivo* motility and invasion without increasing cancer cell proliferation. We conclude that early in cancer evolution NR2F1 serves as a barrier to dissemination of ECCs by repressing a partial EMT program triggered by HER2 signaling. These studies provide novel insight into early changes in cancer progression that may result in early metastatic seeding.

## RESULTS

### NR2F1 downregulation favors a motile phenotype and dissemination in early breast cancer cells

p38α/β signaling positively regulated NR2F1 in HNSCC cells^14,15^. We also found that the phosphorylation of both p38α and p38β and NR2F1 expression were reduced in early lesions of MMTV-HER2 mice when compared to normal mammary glands from FvB female siblings^13,14,16^, suggesting a potential co-regulation. However, whether downregulation of NR2F1 in HER2+ cells was functionally linked to a reduction in p38α/β activity was unclear. To this end, we treated young MMTV-HER2 females at 14-20 weeks of age with a p38α/β inhibitor (SB203580), when this kinase is active in the mammary epithelium^13^. p38α/β inhibition further downregulated *Nr2f1* mRNA and protein levels in mammary glands when compared to vehicle group (**Fig. 1A**). Moreover, *MKK3*^-/-^/*MKK6*^+/-^*mice*, with targeted disruptions of the *MKK3* and *MKK6* genes, both upstream activators of p38, also showed reduced NR2F1 levels compared to wild type (WT) C57BL/6 mice (**Sup. Fig. 1A**) supporting that in the normal mammary gland MKK3/6-p38α/β signaling is required to maintain expression of NR2F1. Accordingly, overexpression of a constitutive active p38α mutant in MMTV-HER2 primary tumor-derived cells (low/negative for phospho-p38^13^ and NR2F1^14^) caused strong upregulation of *Nr2f1* mRNA levels and slightly upregulated protein levels (**Sup. Fig. 1B**).

**Figure 1.**
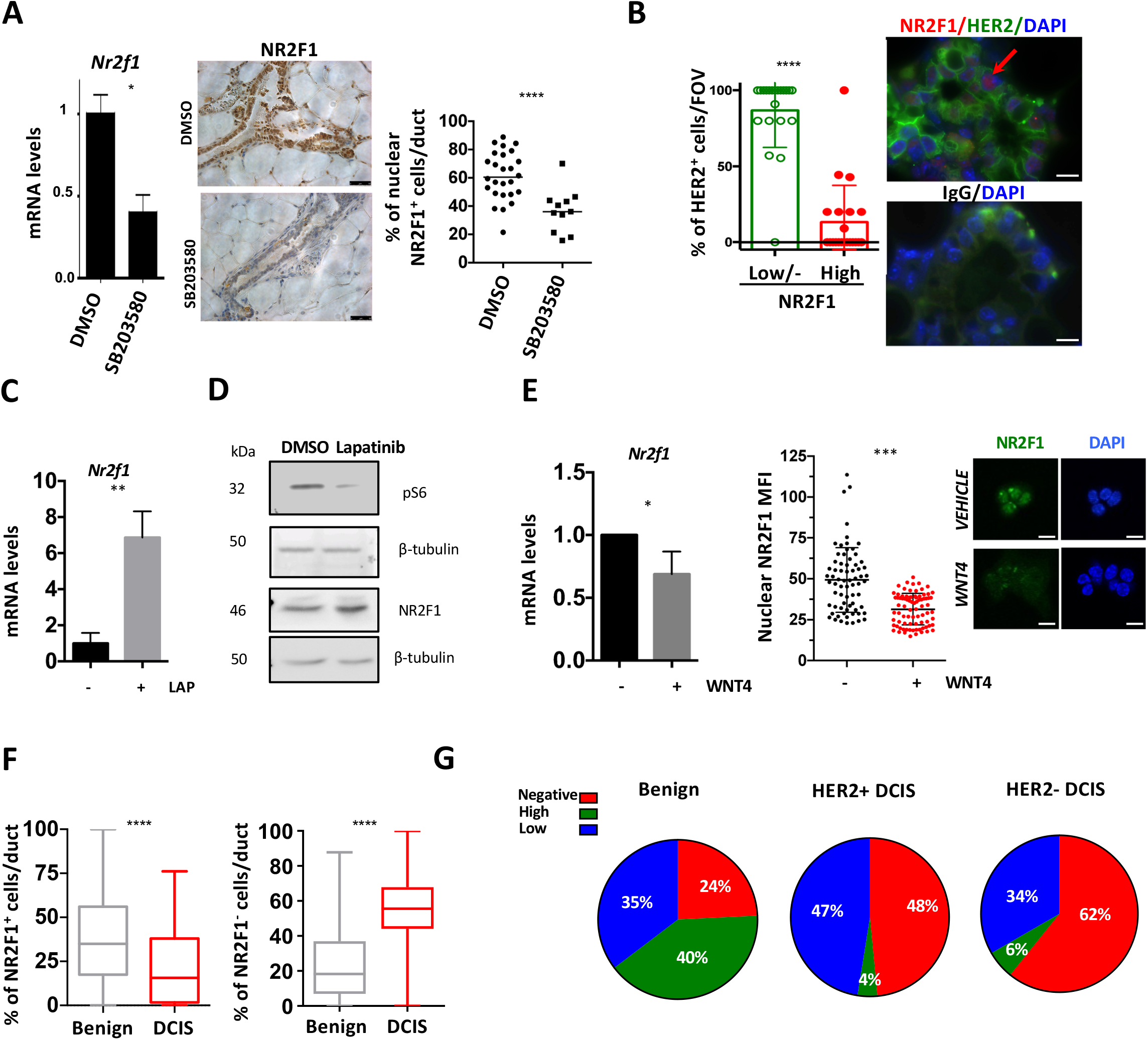
NR2F1 is downstream of p38 and HER2 signaling in MECs. **A. NR2F1 is downstream of p38 signaling in MECs.** *Left panel*: *Nr2f1* mRNA levels in mammary gland tissue obtained from 19-week-old MMTV-HER2 females that were i.p. treated with SB203580 (10 mg/kg every 48 hrs) or vehicle (DMSO) for 2 weeks. *Middle panel*: MMTV-HER2 mammary gland sections from mice treated as above and stained for NR2F1 by immunohistochemistry (IHC). Scale bar, 50 μm. *Right panel*: Graph shows IHC quantification. Number of animals= 2 per condition, number of ducts analyzed, DMSO=26, SB203580=11. Mann-Whitney test. **B. HER2 positive cells display low/negative levels of NR2F1 in MECs.** Mammary glands from 18-week-old MMTV-HER2 females were stained by immunofluorescence (IF) for HER2, NR2F1 and DAPI (Upper panel) or matched IgG isotype (Lower panel). Graph represents the percentage of HER2 positive MECs per field of view (FOV) that is either NR2F1^HIGH^ or NR2F1^LOW/NEGATIVE^. Arrow shows a HER2^+^/NR2F1^HIGH^ cell. Number of animals=2 per condition. Mann-Whitney test. Scale bar, 10 μm. **C. Lapatinib treatment restores *Nr2f1* mRNA levels in early cancer cells.** QPCR for *Nr2f1* mRNA in sphere cultures from MMTV-HER2 early cancer cells (17-week-old females) treated with DMSO or Lapatinib (LAP, 0.5 μM, 24 h). Representative experiment of 3 independent experiments. Student’s unpaired t-test. **D. Lapatinib treatment restores NR2F1 protein levels in primary tumor-derived cells.** Isolated primary tumor cells were treated with lapatinib as in **C** and western blot was performed for the indicated antigens. Representative of 3 independent experiments. **E. WNT4 represses *Nr2f1* expression.** Acini from MMTV-HER2 early cancer cells were treated with WNT4 or vehicle for 24 hours. *Left panel*: QPCR for *Nr2f1*. Three independent experiments. Student’s unpaired t-test. *Right panel*: IF for NR2F1. Scale bar, 10 μm. Graph shows nuclear NR2F1 mean fluorescence intensity (MFI). Student’s unpaired t-test. **F. NR2F1 levels are downregulated in DCIS samples.** DCIS samples (n=13 patients) were stained for NR2F1 by IF (**Sup. Fig. 1D**). Regions with DCIS features were corroborated with the pathologist. Graphs show the percentage of nuclear NR2F1^+(HIGH+LOW)^ (as shown in **Sup Fig. 1D**) and NR2F1^-^ staining in benign adjacent tissues *vs*. DCIS. Mann-Whitney test. **G. NR2F1 levels are downregulated in HER2+ and HER2-DCIS samples.** NR2F1 was stained by IF in HER2- and HER2+ DCIS samples and benign adjacent tissues (as shown in **Sup. Fig. 1D**). Her2 status of DCIS samples was determined by a pathologist after IHC against Her2. Pie graphs show nuclear NR2F1^HIGH^ (green), NR2F1^LOW^ (blue) and NR2F1^negative^ (red) percentages in benign adjacent and DCIS samples.

As inhibition of p38α/β activity resulted in lower expression of NR2F1 in early lesions (**Fig. 1A, Sup. Fig. 1A, B**) and HER2 signaling appeared to inhibit p38α/β activity^13^, we next correlated the levels of expression of NR2F1 in HER2 positive ECCs from MMTV-HER2 females. We observed that around 90% of mammary cells expressing HER2 expressed low/negative levels (based on fluorescence intensity threshold) of NR2F1 protein by immunofluorescence (IF) staining (**Fig. 1B**). Furthermore, inhibition of HER2 (and HER1) oncogene with lapatinib restored NR2F1 levels in ECCs and cells derived from MMTV-HER2 overt primary tumors (**Fig. 1C&D**). We did not detect any changes in *Nr2f1* mRNA levels upon lapatinib treatment in normal FvB mammary epithelial cells (**Sup. Fig. 1C**). In addition, we had previously shown that WNT signaling is key in promoting the early dissemination phenotype^6^. Importantly, we observed downregulation of NR2F1 expression upon treatment of ECCs with WNT4 (**Fig. 1E**), a member of a progesterone-driven signature that mediates dissemination of ECCs^6^. We conclude that the activation of HER2 and WNT4 signaling causes reduced expression of NR2F1 in MMTV-HER2 ECCs.

We next verified whether the loss of NR2F1 observed ECCs in mouse models was also observed in human DCIS. We stained for NR2F1 and observed significant downregulation in the percentage of nuclear NR2F1 positive cells together with an increase in the percentage of NR2F1 negative cells in DCIS when compared to benign adjacent lesions (**Fig. 1F&G, Sup. Fig. 1D**). Then, we correlated NR2F1 levels in DCIS with HER2 status and observed that HER2+ DCIS samples remarkably increase the percentage of NR2F1 negative cells (p<0.0001) while reducing the percentage of NR2F1 positive cells (p<0.0001) when compared to benign adjacent lesions (**Fig. 1G, Sup. Fig. 1E**). Overall, these results indicate that HER2+ DCIS human samples correlate with low/negative NR2F1 levels the same way we showed in the MMTV-HER2 mouse model (**Fig. 1B**). It is however interesting that HER2-DCIS samples were also majority low/negative for NR2F1 when compared to benign adjacent lesions (**Fig. 1G**). In fact, NR2F1 protein levels were also reduced in early lesions from MMTV-Myc females^14^. These results suggest that alternative mechanisms, such as Myc oncogene or progesterone-induced paracrine signals as shown in **Fig. 1E,** could be involved and that the loss of NR2F1 is by itself a crucial step in the early steps of cancer progression.

NR2F1 has been linked mainly to growth suppression, lineage commitment and differentiation^14,17,18^. However, some studies have shown a role in promoting cell migration and axogenesis^19,20^. Further, because reduced p38α/ß signaling in mammary epithelial ECCs caused accelerated early dissemination, we wondered whether NR2F1 might play a role in the migration of MMTV-HER2 ECCs. To this end, we isolated ECCs from 16-18-week-old MMTV-HER2 females or from animals carrying primary tumors (PT) and seeded them in 3D Matrigel to form acini. In ECC acini the percentage of HER2 positive cells per acinus was approximately 80% (**Sup. Fig. 2A**). MMTV-HER2 ECC acini show a higher frequency (80%) of cancer cell invasive phenotype compared to acini formed by late MMTV-HER2 cancer cells from primary tumors (~5%), or normal FvB mammary epithelial cells (~40%) (**Sup. Fig. 2B**). ECCs acini transduced with a lentiviral vector carrying an RFP-reporter for NR2F1 activity were quantified by confocal microscopy. We observed consistent intra-acinar NR2F1-reporter activity. However, only 30% of invading cells had NR2F1 activity **(Fig. 2A)**, suggesting that ~70% of ECCs that invade outwards from the acini have low NR2F1 activity.

**Figure 2.**
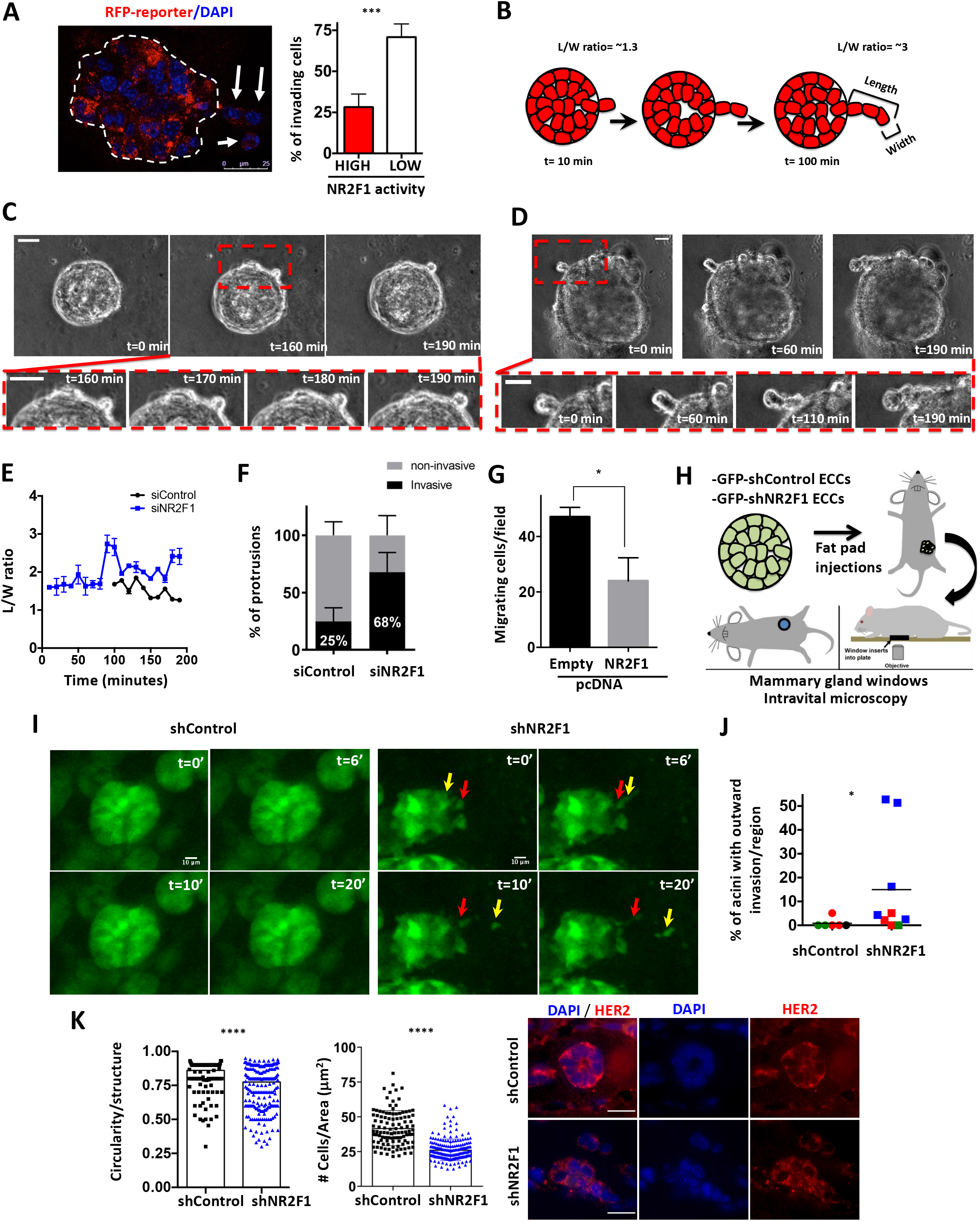
NR2F1 inhibits ECCs invasion. **A. Invading ECCs downregulate NR2F1 activity.** Early cancer cells transduced with RFP reporter plasmid for NR2F1 activity were seeded on Matrigel and fixed. Nuclei were stained with DAPI. Graph shows the quantification of the percentage of invading cells with high or low RFP levels. Arrows show invading cells with low RFP. Student’s unpaired t-test. **B. Schematic of outward invasive protrusions over time.** The length over width of the outward protrusions was measured over time in shControl and shNR2F1 early acini. **C and D. *Nr2f1* KD promotes invasion of ECC into Matrigel.** MMTV-HER2 early cancer acini were transfected with siRNA against *Nr2f1* or control every 24 hours for 2 days. Then, time-lapse imaging was performed for 3 h every 10 minutes. Representative still images from siControl (**C**) and siNR2F1 (**D**) acini taken at the indicated time points are shown. Scale bars=10 μm. **E. *Nr2f1* KD promotes invasion of ECC into Matrigel.** Graph represents the length (L) over width (W) ratios of the protrusions showed in **C** (siControl) or **D** (siNR2F1). Data points represent the mean ± S.D of the measurements taken by 2 independent operators. **F. *Nr2f1* KD induces invasive protrusions over time.** The percentage of protrusions with invasive phenotype (a single elongated cell or cell file columns invading the Matrigel) or non-invasive phenotype was plotted for siControl (n=14) and siNR2F1 (n=14) acini from two independent experiments. **G. NR2F1 overexpression decreases migration of early cancer cells.** Early cancer cells were transfected with either empty or NR2F1 expressing pcDNA vector. Next day, cells were seeded on top of transwell^®^ inserts (8 μm pore size) in serum-free medium. Inserts were added to wells containing complete medium for 16 hours. Then, cells were fixed, stained with DAPI and the number of migrating cells per field was determined. Experiment is representative of 2 independent experiments, 3 technical replicates per condition. Student’s unpaired t-test. **H. High resolution intravital imaging using mammary gland windows.** Schematic of ECCs (early cancer cells) transduced with shControl or shNR2F1 lentiviruses and injected (100,000 cells/site) in the mammary fat pad of nude mice. Ten to sixteen days later mice were anesthetized, and the injected cells were imaged through a mammary gland window. Adapted from Entenberg et al., Methods, 2017^21^ **I. Intravital imaging of early cancer cells in the mammary epithelium.** Time-lapse images were acquired every 2 minutes for an average of 30 minutes. Stills represent ducts imaged at different time points from shControl Movie 5 and shNR2F1 Movie 6 (supplementary data-Movie 5 and 6). Arrows show individual cells moving outward from the acinus. Scale bar=10 μm, t=time. **J. *Nr2f1* KD increases the percentage of invasive early cancer cells *in vivo*.** The number of motile cells from acinar structures was quantified in shControl and shNR2F1 groups. Number of animals analyzed=3/group which are represented by different colors, 1-5 regions per animal. Mann-Whitney test. **K. Analysis of ECC structures *in vivo*.** Mammary gland sections from mice treated as in **Fig. 2H** were stained for HER2 and DAPI (representative images are depicted, scale bar=15 μm). The circularity (left panel) and the number of cells per area (μm) (right panel) of each ECC structure were measured in shControl (n=111) vs. shNR2F1 (n=214) groups (3 mice each group) Student’s unpaired t-test.

To assess whether NR2F1 downregulation observed in outward invading cells was functionally dependent on NR2F1, we knocked-down *Nr2f1* using siRNA in MMTV-HER2 ECCs. *Nr2f1* depletion was efficient (**Sup. Fig. 2C&D**) and did not induced apoptosis (**Sup. Fig. 1E**). We used time-lapse microscopy to track and quantify cell dissemination into 3D Matrigel after NR2F1 downregulation in ECCs acini. We observed that while individual cells moving on the surface of siControl acini largely remain rounded; NR2F1 downregulation induced cell file columns invading the Matrigel (**Fig. 2B-D**, **Movies 1-4**). Analysis of the length over width ratio of the protrusions (one cell or cell file columns) (**Fig. 2B**) showed higher elongation towards the Matrigel (ratios >2.4) in the siNR2F1 group when compared to control group (**Fig. 2E, Supp. Fig. 2F&G**). The number of protrusions per acini did not change between siControl *vs*. siNR2F1 group (**Supp. Fig. 2H**). However, we observed higher percentage of protrusions with invasive phenotype (a single elongated cell or cell file columns invading the Matrigel) in siNR2F1 group when compared to siControl group (**Fig. 2C, D & F**). Moreover, overexpression of NR2F1 (**Sup. Fig. 2I**) in ECCs was able to significantly decrease their migratory capacity, as shown by migration assay in transwell^®^ inserts (**Fig. 2G**).

To more stringently test the function of NR2F1 as a barrier to early dissemination, we quantified how modulation of NR2F1 expression affects the invasion of ECCs *in vivo*. To this end, we performed intravital imaging through mammary fat pad windows^21^ (**Fig. 2H**) of GFP-tagged ECCs stably expressing either shControl or shRNA targeting NR2F1 (**Sup. Fig. 2J**). Intravital time-lapse imaging revealed that GFP positive cells could be readily found viable and organized in acinar-like structures (representing more than 60%) or duct-like structures (~40%) at the same proportions in shControl *vs*. shNR2F1 injected ECCs (**Sup. Fig. 2K**). However, the two-photon microscopy showed that NR2F1 downregulation significantly increased the dissemination of ECC into the surrounding stroma when compared to shControl ECCs (**Fig. 2I** and **J Movies 5, 6**). Analysis of the acini morphology showed less circularity and fewer cells per area in shNR2F1 acini than shControl acini suggesting less compacted structures due perhaps to increased motility (**Fig. 2K**). We conclude that NR2F1 expression limits a program of cell motility, invasion and dissemination in acinar structures of MMTV-HER2 ECCs *in vitro* and *in vivo*.

### NR2F1 blunts the expression of specific EMT regulators in early breast cancer cells

We previously published that ECCs gained expression of the EMT gene TWIST1 among others^13^. We hypothesized that if NR2F1 limits early dissemination, then it might regulate components of the EMT program. In support of our hypothesis, we found that depletion of *Nr2f1* mRNA decreased E-cadherin levels and junctions in early MMTV-HER2 acini at the mRNA and protein levels (**Fig. 3 A&B).** Furthermore, the EMT master regulators *Twist1* and *Zeb1* were upregulated in MMTV-HER2 ECCs acini upon *Nr2f1* knockdown (KD) (**Fig. 3A, C**). Interestingly, we also found that upon *Nr2f1* KD the levels of *Prrx1*, a developmental regulator of EMT involved in metastasis^22^, were upregulated (**Fig. 3A, D**). To determine whether p38 also regulated *Zeb1* and *Prrx1*, we treated ECC acini with either p38 inhibitor SB203580 or DMSO. We observed upregulation of *Zeb1* and *Prrx1* levels in SB203580-treated acini when compared to DMSO-treated acini (**Sup. Fig. 3A**). We also observed a reduction of E-cadherin in FvB normal acini upon *Nr2f1* KD, however it was not accompanied by transcriptional changes of EMT master regulators **(Sup. Fig. 3B)** supporting that the revealed mechanisms are specific to the oncogenic HER2-regulated early progression stages. These results strongly suggest that NR2F1 represses the acquisition of a mesenchymal-like phenotype in luminal early MMTV-HER2 cells before tumor initiation.

**Figure 3.**
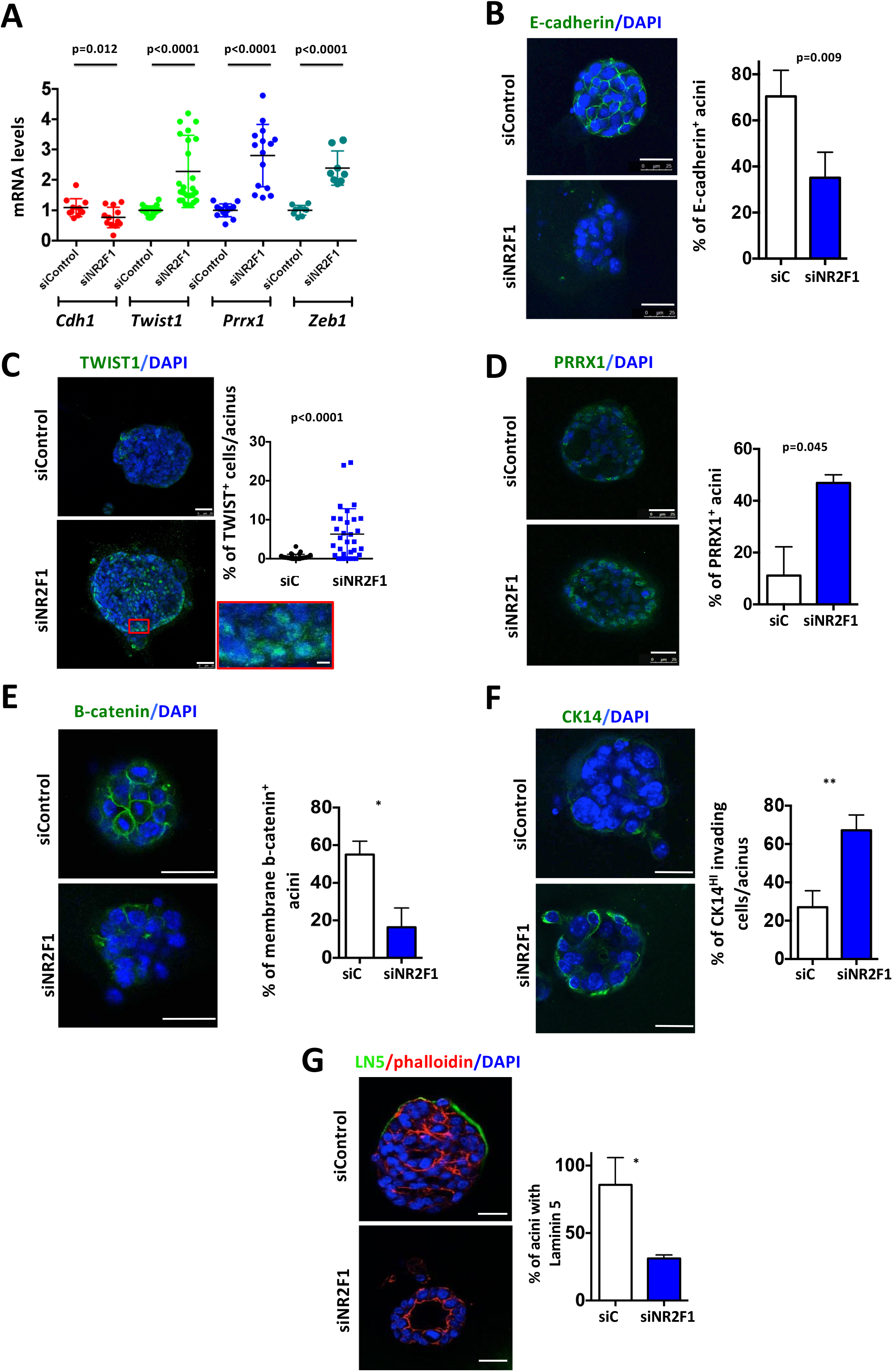
NR2F1 blocks an invasive phenotype. **A. *Nr2f1* KD increases levels of EMT factors in ECCs.** MMTV-HER2 early cancer cells were seeded in Matrigel for 4 days and then cultured acini were transfected with either siControl or siNR2F1 for 48 h. QPCR was performed for the indicated genes. Animals per gene= 3 to 9. Student’s unpaired t-test. **B. *Nr2f1* KD decreases E-cadherin protein levels in ECCs.** Immunofluorescence (IF) staining for E-cadherin in acini obtained as in **A**. Graph shows percentage of acini with E-cadherin positive staining. An acinus is considered positive when containing more than 3 z-stacks with more than 3 positive cells each. Representative images of one out of 3 independent experiments are shown. Acini analyzed/experiment= 40 for siControl; 40 for siNR2F1. Student’s unpaired t-test. **C. *Nr2f1* KD increases TWIST1 protein levels in ECCs.** IF for TWIST1 in acini obtained as in **B**. Graph shows percentage of cells that were positive for TWIST1 per acinus. Experiment is representative of 2 independent experiments. Acini analyzed= 53 for siControl; 52 for siNR2F1. Student’s unpaired t-test. Scale bars, 25 μm and 5 μm (inset). **D. *Nr2f1* KD increases PRRX1 levels in ECCs.** IF for PRRX1 in acini obtained as in **B**. Graph represents percentage of PRRX1 positive acini. Two independent experiments. Acini analyzed= 18 for siControl; 26 for siNR2F1. Student’s unpaired t-test. Scale bars, 25 μm. **E. NR2F1 KD decreases β-catenin membrane localization in ECCs.** IF for total β-catenin in acini obtained as in **B**. Graph shows percentage of acini positive for total β-catenin. An acinus is considered positive when containing more than 5 positive cells with β-catenin localized in the membrane. Two independent experiments. Acini analyzed= 22 for siControl; 28 for siNR2F1. Student’s unpaired t-test. Scale bars, 25 μm. **F. *Nr2f1* KD increases the percentage of CK14 positive invading cells.** IF for CK14 in acini obtained as in **B** (from 14-week-old MMTV-HER2 females). Graph shows percentage of invading cells positive for CK14 per acinus. Two independent experiments. Acini analyzed= 15 for siControl, 26 for siNR2F1. Student’s unpaired t-test. **G. *Nr2f1* KD disrupts the laminin V deposition pattern in ECC acini.** IF for laminin V in acini obtained as in **B.** Graph shows percentage of acini with laminin 5 (LN5) deposition. Two independent experiments. Acini analyzed= 10 for siControl; 15 for siNR2F1. Student’s unpaired t-test. Scale bar, 25 μm.

We also tested the levels of SNAIL and VIMENTIN in early cancer cells upon NR2F1 KD. The mRNA levels for *Snail* were slightly increased upon NR2F1 downregulation (**Sup. Fig. 3C, left panel**). However, nuclear SNAIL protein levels remained the same in siNR2F1-transfected MMTV-HER2 ECCs acini when compared to siControl acini (**Sup. Fig. 3C, right panel**). In addition, the total levels of VIMENTIN were very low in ECCs (**Sup. Fig. 3D, left panel**) and were not affected by NR2F1 KD (**Sup. Fig. 3D, right panel).** These findings support that only certain components of the EMT program are controlled by NR2F1.

Next, we investigated whether NR2F1 controls β-catenin localization as this Wnt-regulated signal was associated with early dissemination in our prior studies^13^. IF staining using total β-catenin antibody showed less membranous localization in NR2F1-depleted acini than in control acini (**Fig. 3E**), similar to what we had observed in ECCs when p38 was inhibited^7^. To map the organoid molecular changes to the *in vivo* scenario we injected the shControl or shNR2F1 ECCs described in **Fig. 2H** into the fat pad of nude mice and then stained for β-catenin. In agreement with our *in vitro* acini assays (**Fig. 3E**), we observed a reduction in the percentage of acini with β-catenin localized to the membrane in shNR2F1 ECCs when compared to shControl group (**Sup. Fig. 3E**). Unfortunately, IF detection of nuclear ß-catenin was not possible in the 3D cultures or *in situ* staining from mammary glands as previously shown due to the limited sensitivity of the technique^13^. Moreover, unlike what we observed with p38α/β inhibition, the mRNA levels of *Axin2*, a transcriptional target of Wnt/β-catenin that creates a negative feedback loop to silence the signaling pathway, did not increase upon NR2F1 KD (**Sup. Fig. 3F**). We conclude that the decreased β-catenin membranous localization might be due to the decreased E-cadherin levels at cell-cell junctions rather than a full activation of the pathway.

It has been previously shown that NR2F1 expression correlates with a more luminal phenotype in mammary epithelial cells^23^ and acquisition of a basal phenotype has been linked to dissemination, metastasis and poor survival^24,25^. Thus, we decided to measure the percentages of invading cells positive for the basal marker CK14. To determine the effect of NR2F1 knock down, we collected ECCs from 14-week-old females, allowed them to form acini, knocked down NR2F1 and measured the percentage of CK14+ invading cells. We observed an increase in the percentage of CK14+ invading cells after NR2F1 knock down (**Fig. 3F**). This data is consistent with an increased CK14 expression in acini-like structures *in vivo*, measured after injection of shNR2F1-ECCs into the fat pad and compared to shControl cells (**Sup. Fig. 3G**). These results suggest that NR2F1 depletion favors a basal phenotype with the acquisition of EMT program regulators in ECCs.

Lastly, we sought to determine the effect of NR2F1 in the invasive phenotype of ECC acini by measuring laminin 5 deposition pattern. We observed that NR2F1 depletion disrupted laminin 5 deposition in MMTV-HER2 ECCs acini (**Fig. 3G**). This result was also obtained when MMTV-HER2 ECCs acini were treated with the p38 inhibitor SB2035807. Based on these results we argue that depletion of NR2F1 in ECCs allows them to acquire a partial EMT program linked to a basal phenotype, which is required for early dissemination.

### Loss of NR2F1 in the mammary epithelium favors early spontaneous dissemination

To assess the tumor-initiating ability of ECCs after NR2F1 KD, we injected either shControl or shNR2F1 ECCs (**Sup. Fig. 2J**) in the mammary fat pad of nude mice and monitored tumor outgrowth. During a follow-up period of 2 months, none of the groups developed tumors (**Fig. 4A**). This is in agreement with our previous work showing that MMTV-HER2 ECCs are largely non-tumorigenic in orthotopic sites^7^. By contrast, we observed tumor take in all mice injected with late MMTV-HER2 tumor cells as previously shown^7^ (**Fig. 4A**). shNR2F1-ECCs injected into the fat pad had the same percentage of cells positive for the proliferative marker phospho-histone 3 (PH3) when compared to shControl cells (**Fig. 4B**). The same result was observed when ECCs were transfected with a different siRNA sequence against NR2F1, cultured in 3D acini and stained for PH3 (**Sup. Fig. 3G**). Notably, the percentage of TWIST1 was higher in shNR2F1-injected ECCs than shControl cells (**Fig. 4C**), confirming our previous results *in vitro* (**Fig. 3C**) and suggesting that a motility and invasion program is active. In order to corroborate our hypothesis concerning the role of NR2F1 in spontaneous dissemination to the lungs, we injected shControl or shNR2F1 ECCs in the mammary fat pad of nude mice and 10-16 days later we stained eDCCs in the lungs (**Fig. 4D**). We observed that the number of lung single-eDCCs was higher in shNR2F1 group when compared to shControl group (**Fig. 4E**) supporting that NR2F1 represses systemic dissemination to distant organs such as the lung. At this time point, there was no difference in the percentage of proliferative eDCCs measured by PH3 staining (**Fig. 4F**) and no metastases were found in either group, as metastases take longer to develop due to a prolonged dormancy phase^13^. Overall, our results suggest that during early stages of cancer progression NR2F1 primarily functions to limit an EMT- and basal-like dissemination program activated by HER2 signaling and linked to motility and invasion process and not to limit a “growth” program (**Fig. 5**).

**Figure 4.**
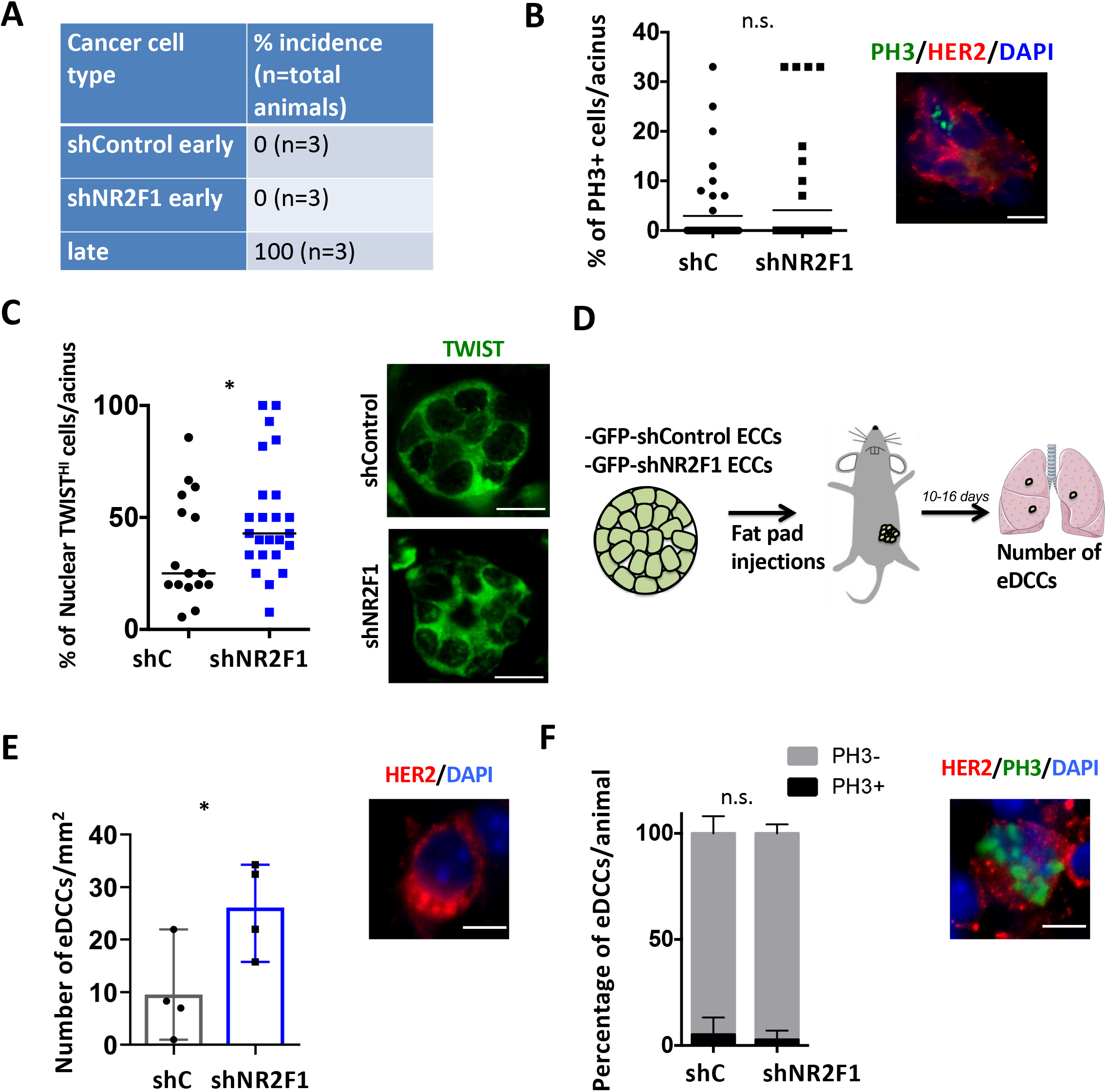
NR2F1 inhibits dissemination of MMTV-HER2 early cancer mammary epithelial cells. **A. NR2F1 KD does not affect the orthotopic tumor take of early cancer cells.** shControl and shNR2F1 early cancer cells grown in organoid media were injected (100,000 cells/site) in the mammary fat pad of nude mice. MMTV-HER2 late cancer cells grown in organoid media were also injected (100,000 cells/site) for comparison purposes. Tumor development was monitored weekly for 2 months and the volume (mm3) of tumors was measured if present. Table represents the percentage of tumor incidence in a period of 60 days. **B. NR2F1 KD does not affect the percentage of PH3 positive cells in the mammary epithelium.** IF staining for PH3 and HER2 in mammary glands that were injected with shControl or shNR2F1 ECCs as in **A** and collected after 10 to 16 days. Graph shows percentage of PH3 positive cells per acinus. Number of mice=2/group. Unpaired student *t*-test. Scale bar=25 μm. **C. NR2F1 KD increases the percentage of early cancer cells with nuclear TWIST1.** Same mammary gland tissues used in **Fig. 4B** were stained with anti-TWIST1 antibody and DAPI. Number of mice=2/group. Scale bar=10 μm. Student’s unpaired *t-* test. **D. Spontaneous dissemination of ECCs.** Schematic showing ECCs transduced with shControl or shNR2F1 lentiviruses and injected (100,000 cells/site) in the mammary fat pad of nude mice. 10-16 days later mice were euthanized and eDCCs were quantified by IF. **E. NR2F1 KD increases the number of single eDCCs in the lung.** shControl and shNR2F1 ECCs were injected in the mammary fat pad of nude mice as described in **Fig. 4D.** After 10-16 days, lungs were sectioned and stained for HER2 to identify DCCs. The graph shows the number of DCCs per lung area. Number of mice=4/group, number of DCCs analyzed= 4,399 for shControl; 4,305 for shNR2F1. Representative image shown, scale bar=5μm. Student’s unpaired t-test. **F. NR2F1 KD does not change the percentage of PH3+ eDCCs in the lung.** Same lungs as in **Fig. 4E** were also stained against PH3, HER2 and DAPI. The graph shows the number of PH3 positive or negative eDCCs. Number of mice=4/group, number of eDCCs analyzed= 4,399 for shControl; 4,305 for shNR2F1. Image depicts a PH3+ eDCCs, scale bar=5μm. Student’s unpaired t-test.

**Figure 5.**
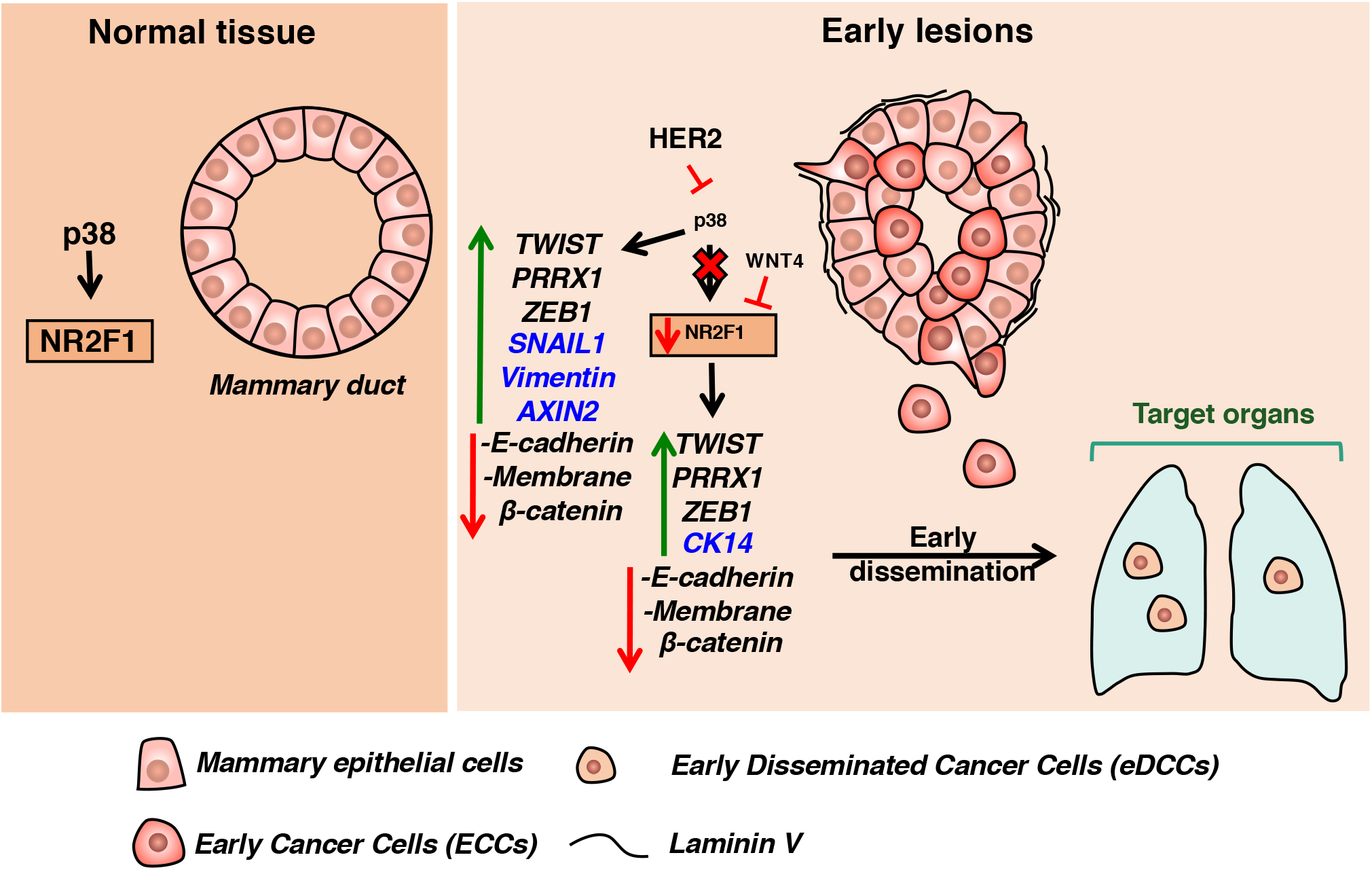
Graphical summary. In normal tissue, mammary epithelial cells regulate normal lumen formation during mammary acinar morphogenesis via p38 activity^13^. Increased HER2 expression reduces p38 activity prompting accumulation of anoikis-resistant epithelial cells and, thus, the formation of early hyperplastic lesions. Based on our new data, p38 inactivity reduces NR2F1 levels in early cancer cells (ECCs) leading to upregulation of EMT markers (e.g. Twist1, Prrx1 and Zeb1), disorganization of Laminin 5, upregulation of cytokeratin 14 (CK14) and reduction of β-catenin membrane localization and the expression of the epithelial marker E-cadherin. The NR2F1-dependent EMT program regulates a sub-set of factors, which does not recapitulate the whole p38-dependent program (factors in blue specific for each program). Moreover, low NR2F1 levels facilitate invasion and dissemination to distant organs. Thus, NR2F1 acts as a barrier for early dissemination and its loss in the mammary epithelium could indicate the presence of early disseminated cancer cells (eDCCs).

## DISCUSSION

Our study shows a novel role for NR2F1 in limiting a motility program that involves features of an EMT and basal mammary epithelium biology that is aberrantly activated downstream of HER2. Our study expands our understanding on how an early-established program in which HER2 primarily deregulates motility and invasion favors dissemination of MMTV-HER2 ECCs to secondary organs before activating an oncogene addictive growth program (**Fig. 5**).

Our findings show that NR2F1 is positively regulated by p38α/ß signaling and is repressed by HER2 and WNT4 pathway in early mammary cancer cells. Supporting the idea of a linear signaling pathway that has p38α/ß upstream of NR2F1 is the fact that both positively control the expression of E-cadherin levels. Interestingly, NR2F1 has been described as a modulator of *Cdh1* mRNA levels in P19 embryonic carcinoma cells^19^. Moreover, NR2F1 blocked the expression of ZEB1 and TWIST1, known negative regulators of E-cadherin^26,27^. This is in agreement with our previously published data, which shows that the pharmacological p38α/ß inhibition decreased E-cadherin mRNA expression and junctions and induced the expression of *Twist1* in premalignant MCF10A cells overexpressing HER2^13^. However, in MCF1A-HER2 cells we had observed transcriptional changes in *Axin2* upon p38 inhibition that were not observed in freshly isolated MMTV-HER2 ECCs upon NR2F1 KD (**Sup. Fig. 3F**). This discrepancy in the regulation of *Axin2* may be due to different cell contexts as MCF10A cells are basal in origin and freshly isolated HER2+ ECCs are mainly derived from the luminal compartment. Alternatively, p38 may be regulating another factor (e.g. ATF2) to control Axin2 expression independently of NR2F1. Further investigation is required to solve this discrepancy.

Simultaneous overexpression of TWIST1 and depletion of E-cadherin in normal mammary epithelial cells has been shown to increase motility of invading cells expressing basal CK14 marker^28^. Our data shows that depletion of NR2F1 in ECCs favored invading cells with CK14 expression. We also observed that the basal phenotype was accompanied by increased motility and upregulation of EMT factors such as TWIST1, ZEB1 and PRRX1 but without affecting VIMENTIN, SNAIL or *Axin2* expression, further supporting a branching downstream of p38 in the regulation of these factors by NR2F1. PRRX1 is particularly interesting, as PRRX1a and PRRX1b isoforms have been involved in tumor progression and they play important roles in metastatic outgrowth and dissemination, respectively^29^. Whether NR2F1 can modulate the alternative splicing of PRRX1 in the mammary epithelium remains to be elucidated.

NR2F1 has been described as a regulator of tumor dormancy in a variety of tumor types^14^. In fact, NR2F1 expression is relatively low in breast, HNSCC and prostate cancer. However, once cancer cells disseminate to secondary organs and become late DCCs, they regain NR2F1 expression^14,30^. We also showed that NR2F1, downstream of p38 signaling, regulated dormancy of late DCCs in advanced tumor models^14^ and that detection of NR2F1 in DCCs from breast and prostate cancer patient’s bone marrow aspirates correlated with good prognosis^31,32^. Moreover, we showed that the dormancy program of eDCCs was p38-independent^7^. These results indicate that different dormancy programs are present in early *vs*. late DCCs. Whether NR2F1 expression is regained in eDCCs and whether it plays a role in dormancy of eDCCs remains to be elucidated.

Overall, we describe a new function of NR2F1 as a metastasis-suppressor that prevents dissemination of early breast cancer cells. The downregulation of NR2F1 levels has been associated with the deletion of a susceptibility locus in a rat-human comparative genetic analysis^23^. Since early spread has already occurred at the time the patient is diagnosed, determining the association of NR2F1 with these susceptibility loci may predict patients at risk, especially those with family history. Early dissemination might also explain metastatic disease in patients diagnosed with cancer of unknown primary (CUP), which accounts for 3-5% of all solid malignancies^33^. Here, ECCs disseminate from undetectable tumors and may be responsible for the metastatic growth. Moreover, early dissemination events could be the first step in the progression of detectable invasive tumors and contribute to metastasis formation^7,8^. Furthermore, we published that invasive primary tumors may recapitulate some of the early mechanisms of dissemination^6^ and then, NR2F1 loss could also be responsible for dissemination in advanced tumor cells. As a matter of fact, ER+ breast primary tumors with high NR2F1 levels that have been associated with higher metastasis-free proportions could be explained by having less disseminated tumor cells^34^. Our findings support a scenario in which NR2F1, downstream of MKK3/6-p38-MAPK signaling, antagonizes HER2 signaling in the mammary epithelium preventing early dissemination (**Fig. 5**). Interestingly, loss of p38 activity^35,36^ in mammary epithelial cells predisposed these cells to HER2-mediated tumorigenesis. Moreover, inhibition of p38α/ß by SB203580 treatment induced the formation of DCIS-like lesions in young MMTV-HER2 females^16^. By contrast, we observed that NR2F1 KD only inhibited dissemination without affecting proliferation of ECCs, suggesting that the NR2F1-dependent signaling is a specific dissemination sub-program of the p38α/ß pathway. Thus, our findings argue for the existence of a coordinated program involving tumor suppression (p38α/ß-dependent) and inhibition of dissemination (p38α/β- and NR2F1-dependent) at early stages of cancer progression. Thus, the development of therapies aiming to maintain the NR2F1 signal should prevent and/or reduce early dissemination of cancer cells and even metastatic outgrowth.

### Experimental procedures

#### Animals

Animal procedures were approved by the Institutional Animal Care and Use Committee (IACUC) of Icahn School of Medicine at Mount Sinai protocol 2017-0162. C57BL/6J *MKK3^-/-^/MKK6^+/-^* and FvB MMTV-HER2 mice were previously described^37,7^. FVB/N-Tg(MMTVneu)202Mul/J, nude (Foxn1^nu^), wild type C57BL/6J and FVB/NJ animals were purchased from Jackson Laboratories.

#### Isolation of mammary epithelial cells

MMTV-HER2 mice: When processed for isolation of early cancer cells (ECCs), absence of any visible small lesions or palpable tumors was confirmed among all 5 pairs of mammary glands in mice of 14-18 weeks of age. MMTV-HER2 mice were euthanized using CO2 inhalation at 14-18 weeks of age or when overt primary tumors (PT) had formed. Whole mammary glands were minced and digested in Collagenase/BSA at 37°C for 45-60min. Red blood cell lysis buffer was used to remove blood cells and cells were then plated for 10-15min in DMEM+10%FBS in 100mm dishes at 37°C for fibroblast removal. Cells were then incubated in 1mM PBS-EDTA for 15min at 37°C and passed through a 25-gauge needle. Cell suspensions were filtered through a 70 μM filter before counting. For generation of mammosphere cultures, isolated cells were seeded in 6-well ultra-low adhesion plates (Corning #3471) at a density of 1×10^6^ cells per well in 1.5 ml mammosphere medium [DMEM/F12 (Gibco #11320-082), 1:50 B27 Supplement (Gibco #17504-044), EGF (10 ng/ml; Peprotech #AF-100-15-A), 1:100 Pen/Strep (Invitrogen #15070-06)]. The following day another 0.5 ml of mammosphere medium was added to each well. Mammosphere cultures were kept for 7 to 10 days at 37°C before using them for experiments. For the generation of acini, 50,000 isolated cells were seeded on a layer of growth factor-reduced Matrigel (Corning #344230) in assay medium with 2% of Matrigel (Corning #356231). Assay medium [DMEM/F12 (Gibco #11320-082), 2% Horse Serum (Invitrogen #16050-122), 0.5 μg/ml Hydrocortisone (Sigma-Aldrich #H-0888), 100 ng/ml Cholera Toxin (Sigma-Aldrich #C-8052), 10 μg/ml Insulin (ThermoFisher #12585-014), 1:100 Pen/Strep (Invitrogen #15070-06)] was supplemented with 5 ng/ml EGF (Peprotech #AF-100-15-A) immediately before seeding cells. For generation of organoids, 100,000 cells were seeded in a six-well low attachment plate in 2 mL per well of organoid media [DMEM/F12 (Gibco #11320-082), 5% Fetal Bovine Serum (Gibco #10438-026), 20 ng/ml bFGF (R&D Systems (R&D) #233-FB-10), 10 ng/ml EGF (Peprotech #AF-100-15-A), 2.5 μM Rock inhibitor (EMD Millipore #420220), 4 ug/ml Heparin (Sigma-Aldrich #H3393-10KU)] with 3% Matrigel. After every 3-4 days 0.5 ml fresh media with Matrigel was added until used for experimental purposes. For generation of stable cells, 200,000 isolated ECCs were transduced with shControl or shNR2F1 lentiviruses and selected with 1 μg/ml puromycin (Alpha Aesar #J67236) for 10 days.

#### Treatment of acini and mammosphere cultures

After 5-7 days in culture, acini were transfected with siRNA against NR2F1 or control for 48h. Acini were then fixed with 4%PFA and immunofluorescence assays were performed using the indicated antibodies. To measure mRNA changes, acini were transfected with the indicated siRNA and 48h later RNA was extracted from cultures using 1mL Trizol (Invitrogen #15596018) followed by RNA extraction. In some cases, cultures were treated for 24 hours with 1μM Lapatinib (SelleckChem #S2111), 200 ng/ml Wnt4 (R&D #475-WN-005/CF), 5μM SB203580 (Selleck #S1076), or vehicle and RNA was extracted. In some cases, mammospheres growing in suspension conditions were transfected with siRNA against NR2F1 or control for 48h and then RNA was extracted. For evaluation of invasion in response to NR2F1 expression, ECCs were transduced with lentiviral particles carrying an NR2F1 responsive promoter driving RFP expression. The lentiviral vector used, (pLV[Exp]-Puro-COUP-TF cis element>mRFP1), was custom-designed by VectorBuilder (vector ID: VB170830-1011esp) and packaged in-house. Transduced acini were evaluated by Confocal Microscopy using a Leica SP5 confocal microscope and the LAS X Leica imaging software.

#### Live Cell microscopy

3D acini cultures were time-lapse imaged using an inverted microscope Olympus IX-70 with Live Cell enclosed chamber. We recorded 4 positions per condition in parallel for 2 hours with 10 minutes interval. Temperature was maintained at 37°C and CO2 at 5% throughout the imaging session. Movies were made by using either Metamorph^®^ (Molecular Devices) or the open source imaging software Image J^38^. This was done using the services of the Microscopy CoRE at Icahn School of Medicine at Mount Sinai.

#### *In vivo* experiments

Single cell suspension from organoids or mammosphere cultures were spun down at 230g for 4min and then 100,000 cells were suspended in 150μl PBS with 1mM calcium and 0.5mM magnesium (PBS++). Matrigel was then added in a 1:1 ratio. Cells were injected into one inguinal (#4) mammary gland fat pad using a 27-gauge needle. Injection sites were monitored for development of tumors weekly for 2 months and final tumor incidence was calculated. Mice injected with PT derived cells were sacrificed when tumors reached 500 mm^3^ according to IACUC regulations.

For 2-photon intra-vital microscopy of mammary glands, GFP-tagged stable ECCs expressing either shControl or shRNA targeting NR2F1 cells were orthotopically injected. Mice were anesthetized using ketamine/xylazine and a skin flap surgery performed exposing the 4^th^ and 5^th^ mammary glands as previously described^21^. Intravital imaging was performed using an Olympus FV1000 MPE two-laser multiphoton microscope. An Infra-red tunable laser was used at 880nm excitation for imaging of GFP tagged cells. Imaging was done with a 25x 1.05NA (XLPL25XWMP2, Olympus) water immersion objective. For each mouse, fields were imaged at 5μm z steps to a depth of ~50μm, with each stack taken approximately every 2 min for 1 hour. Images were reconstructed and analyzed in ImageJ^38^ utilizing the custom written ImageJ plugin ROI_Tracker^39^. Any residual x-y drift not eliminated by the fixturing window was removed with post processing using the StackReg plugin (https://doi.org/10.1109/83.650848) for ImageJ. All procedures were conducted in accordance with the National Institutes of Health regulations and approved by the Icahn School of Medicine at Mount Sinai animal use committee.

#### Immunofluorescence (IF)

Three-Dimensional cultures were fixed with 4% PFA for 20min at room temperature. Staining was performed as previously described for MMTV-HER2 3D cultures^7^. Briefly, cells were permeabilized using 0.1% Triton X-100 for 20min and blocked using 1x IF wash buffer (components) + 10% Normal Goat Serum (Gibco, #PCN5000) for one hour. Primary antibodies were all used at 1:100 concentration unless specified otherwise. Antibodies used were: E-cadherin (BD Biosciences #610181); beta-catenin (BD biosciences 610153); Laminin V (Progen 10765); CK14 (Biolegend #905301); 1:25 NR2F1 (Abcam #ab181137); PRRX1 (Novus Biologicals #NBP2-13816); TWIST (Sigma-Aldrich#ABD29); 1:50 HER2 (R&D #OP15L); PH3 (Cell Signaling #9701S). The following secondary antibodies were used at 1:1000 dilution: Alexa fluor-488 goat anti-mouse (Invitrogen #A-11001) and Alexa fluor-568 goat antirabbit (Invitrogen #A-11036). F-Actin was detected using Alexa Fluor 568 Phalloidin (Invitrogen #A-12380). Chambers were removed from slides and wells were fixed and mounted with ProLong™ Gold Antifade reagent with DAPI (Invitrogen P36931). Acini were imaged in 3D using a Leica SP5 confocal microscope and the LAS X Leica imaging software. Mammary gland section imaging was done using Leica DMi8 fluorescence microscope using LAS X Leica software. For IF staining of paraffin-embedded tissue: sections were routinely deparaffinized and rehydrated. Antigen retrieval was done in 10mM Sodium Citrate Buffer (pH 6) to later permeabilize using 0.1% Triton x-100 in PBS. Blocking was done using 3% BSA in PBS with 1.5% Normal Goat Serum or Normal Donkey Serum (Sigma-Aldrich #D9663). Primary antibodies at 1:100 dilution – unless otherwise specified – were assessed overnight at 4°C and secondary antibodies at 1:1000 dilution were assessed for one hour at room temperature. The following primary antibodies were used; beta-catenin (Cell Signaling #8480S or BD biosciences #610153); CK14 (Biolegend #905301); 1:50 HER2 (Abcam, #ab2428 or R&D Systems #OP15L); PH3 (Cell Signaling #9701S); TWIST1 (EMD Millipore # ABD29); 1:25 NR2F1 (Abcam #ab181137). The following secondary antibodies were used: Alexa Fluor-488 goat anti-mouse, Alexa Fluor-568 goat antirabbit, Alexa fluor-488 donkey anti-goat, Alexa fluor-647 donkey anti-rabbit (Invitrogen, #A-31573). Slides were mounted using Prolong™ Gold Antifade mounting media with DAPI (Molecular Probes #P36931).

#### Western Blot

Samples were collected in RIPA buffer and centrifuged at 4°C, 18,000 g to clarify lysate. Protein concentration was determined by Pierce^™^ BCA Protein Assay Kit (Thermo Scientific #23225) and a standard BSA curve. Samples were then boiled for 5 min at 95°C in sample buffer (0.04M Tris-HCL pH 6.8, 1%SDS, 1% β-mercaptoethanol and 10% glycerol). SDS-PAGE 10% gels were run in Running Buffer (25mM Tris, 190mM glycine, 0.1% SDS) and transferred to PVDF membranes in Transfer Buffer (25mM Tris base, 190mM glycine, 20% methanol). Membranes were then blocked in 5% milk in TBST. Primary antibodies were left overnight at 4°C. Following washing with TBST buffer HRP conjugated secondary antibodies were left at room temperature for one hour. Western blot development was done using Amersham ECL Western Blot Detection (GE #RPN 2106) and GE ImageQuant LAS 4010. Primary antibodies used were: p38 1:500 (BD Biosciences #612169); GAPDH (Calbiochem #CB1001); NR2F1 1:500: (Abcam #ab181137). Secondary antibodies used were Peroxidase Horse Anti-Mouse IgG (1:5000) (Vector #PI-2000) or anti-rabbit IgG (1:5000) ((Vector #PI-1000).

#### QPCR

RNA was extracted from 2D or 3D cell cultures using Trizol following provider’s recommendation. For early mammary gland tissue RNA was extracted using Qiagen’s RNeasy Lipid Tissue Midi Kit (Qiagen #74804). RNA (1-2 μg) was retrotranscribed into cDNA using RevertAid First Strand cDNA Synthesis Kit (Thermo Scientific # K1621). Quantitative real time-PCR was performed using PerfeCTa SYBR green (Quantabio, #P/N 84069) in Biorad thermocycler. *Gapdh* was used as reference gene control for all plates. Mouse Q-PCR primers *wereGapdh* forward primer 5’-AACTTTGGCATTGTGGAAGGGCTC-3’; *Gapdh* reverse primer 5’-TGGAAGAGTGGGAGTTGCTGTTGA-3’; *Twist1* forward primer 5’-AACTGGCCTGCAAAATCATA −3’; *Twist1* reverse primer 5’-ACACCGGATCTATTTGCATT-3’; *Zeb1* forward primer 5-ACCCCTTCAAGAACCGCTTT-3’; *Zeb1* reverse primer 5-CAATTGGCCACCACTGCTAA-3’; *Nr2f1* forward primer 5’-CCAATACTGCCGCCTCAA-3’; *Nr2f1* reverse primer 5’-GGTTGGAGGCATTCTTCCTC-3’; *Prrx1* forward primer 5’-GCACAAGCAGACGAAAGTGT-3’; *Prrx1* reverse primer 5’-CTGGTCATTGTCCTGCTGAG-3’; *Snai1* forward primer 5’-GGCGGAAGCCCAACTATAGC-3’, *Snai1* reverse primer 5’-AGGGCTGCTGGAAGGTGAA-3’; *Vim* forward primer 5’-AGGAGGCCGAAAGCACCCTGC-3’, *Vim* reverse primer 5’-CCGTTCAAGGTCAAGACGTGCCA-3’; *Cdh1* forward primer 5’-ACAGACCCCACGACCAATGA-3’, *Cdh1* reverse primer 5’-CCTCGTTCTCCACTCTCACA-3’; *Axin2* forward primer 5’-CAAAAGCCACCCAAAGGCTC −3’, *Axin2* reverse primer 5’-TGCATTCCGTTTTGGCAAGG-3’.

#### Patient samples

Paraffin embedded sections from DCIS and invasive breast cancer patient tumors were obtained from the Cancer Biorepository at Icahn School of Medicine at Mount Sinai, New York, NY. Samples were de-identified and obtained with Institutional Review Board approval, which indicated that this work does not meet the definition of human subject research according to the 45 CFR 46 and the Office of Human Subject Research. IF and immunohistochemistry (IHC) analysis was done using samples from 13 DCIS patients.

### HER2 immunohistochemistry and scoring

Sections were routinely deparaffinized and rehydrated before staining procedures. Sections were treated with stabilized 3 % hydrogen peroxide solution for 30 min to block endogenous peroxidase activity, incubated in a water bath immersed in sodium citrate buffer (pH 6.0) for 40 min at 97 °C for antigen retrieval, and incubated with 3 % BSA in PBS for 1 hour to block nonspecific binding, washing thoroughly in PBS between steps. HER2 immunohistochemistry was performed with Anti-c-ErbB2/c-Neu (Ab-3) Mouse mAb (3B5) 1/200 (Sigma-Aldrich t# OP15) overnight at 4°C. After washing, secondary Horse Anti-Mouse IgG Antibody (H+L), Peroxidase (Vectort# PI-2000) was added diluted 1/200 for 1 h at room temperature. Signal was developed for 10 min in DAB solution (Vector #SK-4100 prepared according to manufacturer instructions), sections were then counterstained for 1 min with Harris hematoxylin, rehydrated, and mounted. Negative controls using mouse IgG showed no immunoreactivity.

Evaluation of immunostainings was performed by a trained pathologist and scored according to ASCO/CAP guidelines^40^: negative for 0 (no membrane staining) and 1+ (faint or barely perceptible incomplete membrane staining); equivocal for 2+ (moderate circumferential staining in >10% of tumor cells or strong circumferential membranous staining in ≤10% of tumor cells) and positive for 3+ strong circumferential membranous staining in >10% of tumor cells).

#### Statistical Analysis

Statistical Analysis was done using Prism Software. Differences were considered significant if *P* values were <0.05. For the majority of cell culture experiments, twotailed Student’s t-test was performed unless otherwise specified. For mouse experiments one tailed Mann-Whitney test was used. Sample size chosen was done empirically.

## Supporting information

Supplementary Movie 1 - shControl

Supplementary Movie 2 - shControl

Supplementary Movie 3 - shNR2F1

Supplementary Movie 4 - shNR2F1

Supplementary Movie 5 - shControl

Supplementary Movie 6 - shNR2F1

## Acknowledgements

We want to thank Roger Davis for *MKK3^-/-^/MKK6^+/^-* samples and his valuable feedback. Grant support: CCR17483357 (M.S.S.); MRF-CDA (M.S.S); K22CA201054 (M.S.S); CSBC Pilot Project-Sage Bionetworks (M.S.S.); Schneider-Lesser Foundation Fellow Award (M.S.S); MRA (M.S.S); CCR18547848 (J.J.B-C); K22CA196750 (J.J.B-C); P30-CA196521(J.J.B-C).

## Author Contributions

C.R-T. designed, performed *in vitro* and *in vivo* experiments, collected microscopy data, analyzed data and co-edited the manuscript; N.K. performed experiments and analyzed data; M.J.C. performed transwell migration experiments, IHC staining, analyzed data and co-edited the manuscript; N.S. performed experiments and analyzed data; J.J.B-C. designed and executed intravital imaging; M.A. analyzed IHC staining for HER2 score and identification of DCIS lesions and benign adjacent tissues; J.J. performed statistical analysis; M.S.S. designed and executed intravital imaging, performed *in vitro* and *in vivo* experiments, provided general oversight, collected microscopy data, analyzed data and wrote the manuscript.

## Competing Interests statement

Not applicable.

---

**Supplementary Figure 1.**
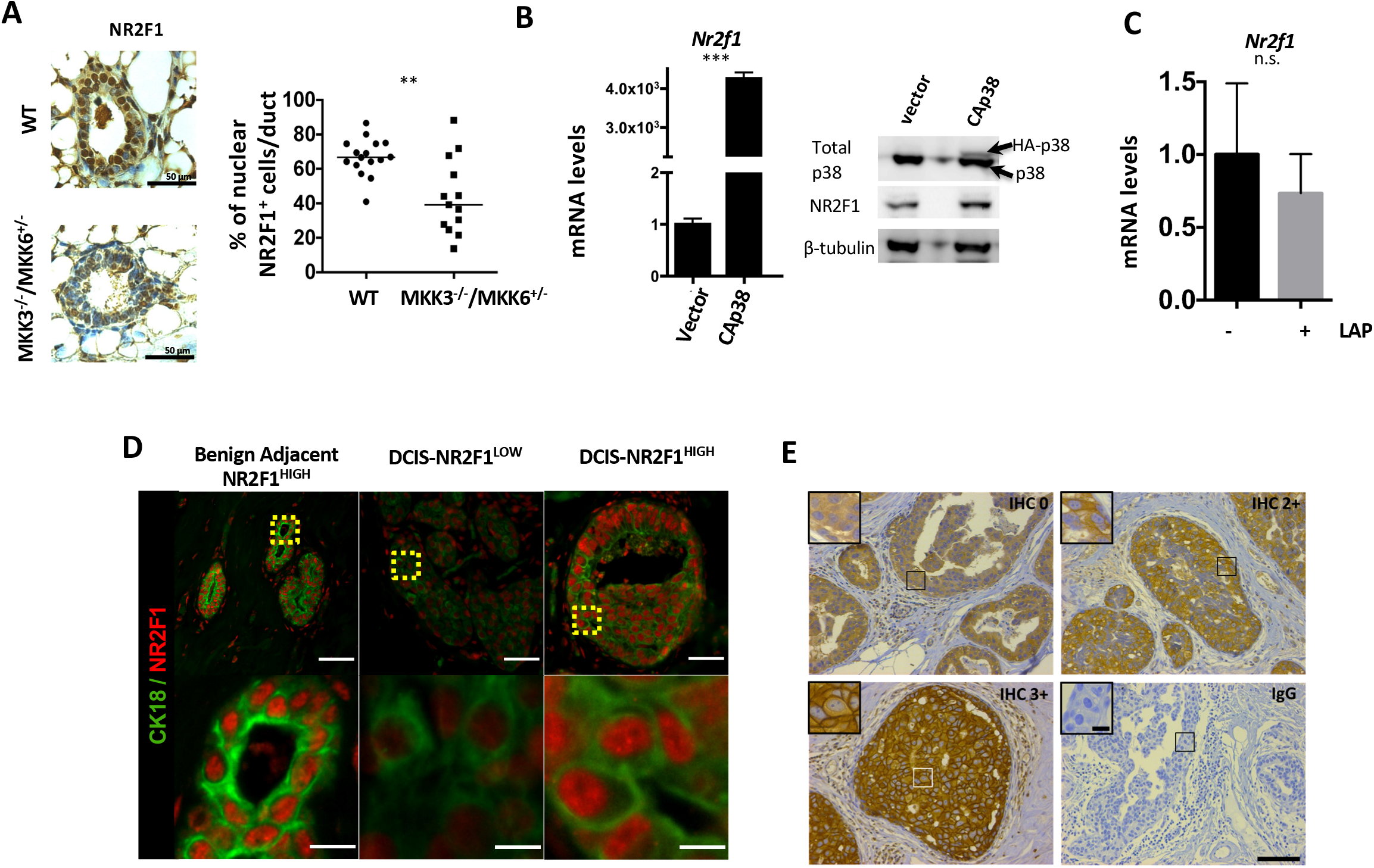
NR2F1 is downstream of HER2 and p38 signals. **A. MKK3^-/-^/MKK6^+/-^ mice display low levels of NR2F1 in mammary epithelial cells.** Mammary gland sections from wild type (WT) C57BL/6J or MKK3^-/-^/MKK6^+/-^ mice were stained for NR2F1 by immunohistochemistry. Graph shows percentage of cells with nuclear NR2F1 staining (NR2F1^+^) per duct. Number of mice=2/group, number of ducts analyzed, WT=25, MKK3^-/-^/MKK6^+/-^=23. Mann-Whitney test. **B. Exogenous expression of the constitutive active mutant p38 (CAp38) increases NR2F1 levels in HER2+ primary tumor-derived cells.** Isolated primary tumor cells were transfected with vector or hemagglutinin (HA)-tagged CAp38 plasmid for 48 hours. CAp38 protein expression was confirmed by western blot and NR2F1 was detected by QPCR and western blot. Student’s unpaired t-test. **C. Lapatinib treatment has no effect on *Nr2f1* expression in FvB mammary epithelial cells.** QPCR for *Nr2f1* mRNA in sphere cultures from FvB-derived mammary cells treated with DMSO or Lapatinib (LAP, 0.5 μM, 24 h). Student’s unpaired t-test. **D. NR2F1 staining in DCIS samples.** Images show immunofluorescence (IF) staining for NR2F1 (red) and Cytokeratin18 (Green) in benign adjacent tissue and DCIS samples. Representative images for NR2F1^HIGH^ and NR2F1^LOW^ staining are shown. Scale bar=50μm. The insert shows a higher magnification of the squared areas. Scale bar=10μm. **E. HER2 staining in DCIS samples.** DCIS HER2 immunohistochemistry (IHC) with score of 0 (IHC 0): no staining or membranous staining that is faint/barely perceptible in <10% of cells, equivocal (IHC 2+): weak to moderate complete membrane staining in >10% of cells, and positive (IHC 3+) scores; and isotype control (IgG). Scale bar=100μM. The insert shows a higher magnification of the squared areas. Scale bar=20μM.

**Supplementary Figure 2.**
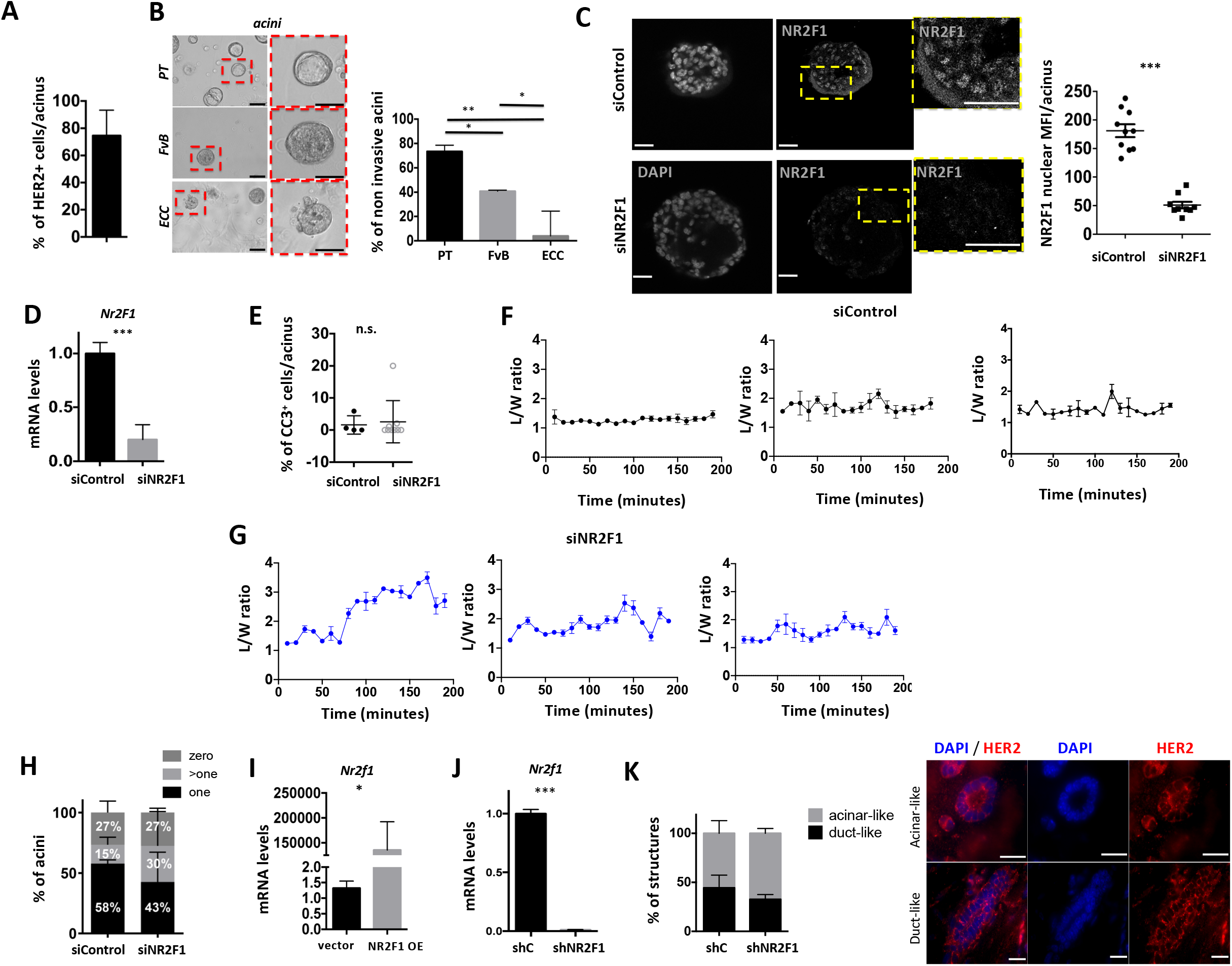
NR2F1 inhibits ECCs invasion. **A. The majority of the cells in cultured acini from MMTV-HER2 mammary epithelial cells are HER2 positive.** Quantification of immunofluorescence (IF) staining for HER2 in MMTV-HER2 early cancer acini (18-week-old female) as percentage of HER2 positive (HER2+) cells per acinus. Approximately 130 cells/acinus were scored. **B. MMTV-HER2 early cancer acini show an invasive phenotype.** Cells derived from MMTV-HER2 primary tumors (PT), FvB mammary glands and early MMTV-HER2 lesions were cultured in Matrigel. After 5 days, brightfield pictures of the cultured acini were taken for each group. Graph shows percentage of non-invasive acini per group. Acini analyzed in PT, FvB and early cancer group=33, 17 and 19, respectively. Student’s unpaired *t*-test. Scale bar = 100μm. **C & D. NR2F1 knockdown confirmation in early cancer MMTV-HER2 acini.** MMTV-HER2 early cancer cells were seeded in Matrigel for 4 days and then cultured acini were transfected with either siControl or siNR2F1 for 48 h. NR2F1 protein levels were detected by IF (**C**). Graph shows average fluorescence intensity in the nucleus. Each dot represents one acinus. Scale bar=25μm. *Nr2f1* mRNA was detected by QPCR (**D**). Student’s unpaired t-test. **E. *Nr2f1* KD in early cancer acini does not affect apoptosis.** Graph shows percentage of apoptotic cells per acinus quantified from IF for cleaved caspase-3 (CC3) in acini treated as in **C**. Student’s unpaired t-test. **F & G. NR2F1 KD promotes invasion in early cancer acini.** Graphs show the length (L) over width (W) ratios of one protrusion/acini in either siControl (**F**) or siNR2F1 (**G**) acini over time (minutes). **H. Acini phenotype proportions in siControl *vs*. siNR2F1 groups.** The percentage of acini with zero, one or more than one protrusion was calculated from the analysis of time-lapse imaging (as shown in **Fig. 2 C and D**) in siControl (n=14 acini) vs. siNR2F1 (n=14 acini) acini from two independent experiments. **I. Confirmation of *Nr2f1* overexpression in early cancer cells.** Early cancer cells were transfected with empty vector or NR2F1 plasmid for 48h and *Nr2f1* expression was measured by QPCR. Student’s unpaired t-test. **J. Generation of shControl and shNR2F1 early cancer cells.** Early cancer cells were transduced with either shControl (shC) or shNR2F1 lentiviruses. Cells were selected with puromycin (1 μg/ml) for 10 days and RNA was extracted to measure *Nr2f1* mRNA levels by QPCR. *Nr2f1* levels were normalized against *Gapdh*. Student’s unpaired *t-* **K. Classification of mammary structures in shControl *vs*. shNR2F1 groups.** ECCs structures obtained from mammary gland sections from mice treated as in **Fig. 2H** were stained for HER2 by IF and DAPI. The percentages of acinar-like and duct-like structures were measured (representative images are depicted, scale bar=15μm). Three mice per group were analyzed.

**Supplementary Figure 3.**
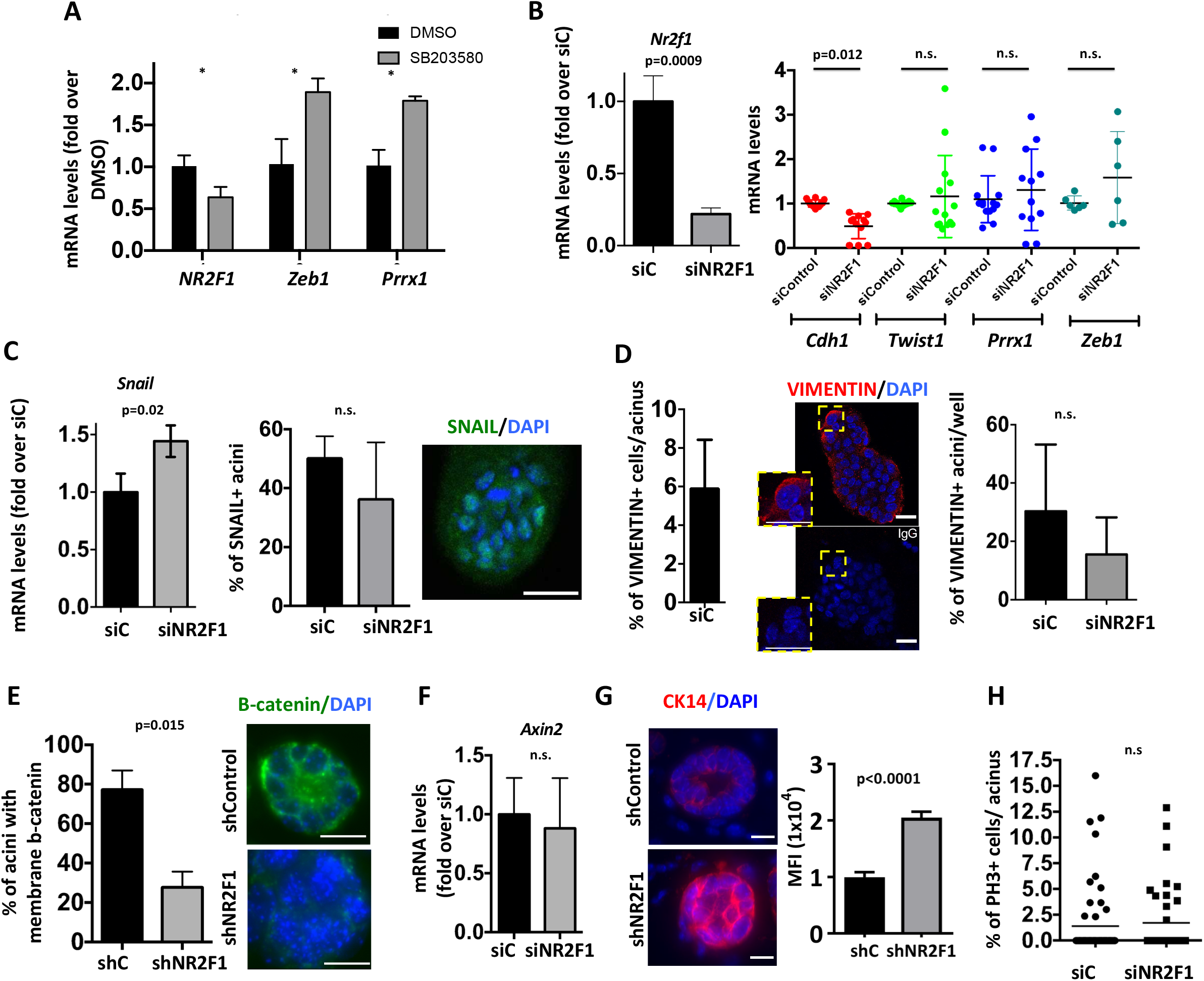
NR2F1 blocks an invasive phenotype. **A. P38 inhibition induces *Zeb1* and *Prrx1* while repressing *Nr2f1* in ECCs.** MMTV-HER2 ECCs isolated from 16-week-old females were grown in acini and treated with either DMSO or SB203580 (5 μM) for 48 hours. RNA was collected and QPCR for the indicated genes was performed. One experiment, 3 replicates per gene. Student’s unpaired t-test. **B. *Nr2f1* KD in FvB-derived MECs reduces E-cadherin expression but does not affect EMT transcription factors.** FvB-derived MECs were seeded in Matrigel for 4 days and then cultured acini were transfected with siRNA against NR2F1 or control for 48 h. KD was validated by QPCR (*Left panel*). QPCR was performed for the indicated genes (*Right panel*). Number of mice per gene= 2 to 6. Student’s unpaired t-test. **C. *Nr2f1* KD does not affect SNAIL protein levels but marginally induces RNA levels.** MMTV-HER2 ECCs isolated from 16-week-old females were grown in acini and fixed after 48 h of siControl or siNR2F1 transfection. Acini were stained with anti-SNAIL antibody. Graph shows average of percentage of SNAIL positive acini per well. An acinus is considered positive when containing more than 3 z-stacks with more than 1 cell positive for nuclear SNAIL each. Two independent experiments. Acini analyzed= 33 for siControl; 35 for siNR2F1. Student’s unpaired t-test. Scale bar=35 μm. **D. *Nr2f1* KD does not affect the low VIMENTIN protein levels.** Acini obtained as in **B,** were stained with anti-VIMENTIN antibody. A representative IF image of the siControl group is shown. *Left panel*= graph shows percentage of VIMENTIN positive cells per acinus. *Right panel*= graph shows percentage of VIMENTIN positive acini per well in siControl and siNR2F1 conditions. An acinus is considered positive when containing more than 3 z-stacks with one cell positive for VIMENTIN. Representative result of 2 independent experiments. Student’s unpaired t-test. Acini analyzed=33 for siControl; 35 for siNR2F1. Scale bars=20 μm. **E. *Nr2f1* KD decreases the percentage of acini positive for membrane β-catenin.** Same mammary gland tissues used in **Fig. 4B** were stained with anti-β-catenin antibody and DAPI. Graph shows percentage of acini with membranous β-catenin localization. Number of mice=2/group. Acini analyzed=67 for shControl; 79 for shNR2F1. Scale bar=25 μm. **F. *Nr2f1* KD does not affect Axin2 expression in ECCs.** QPCR for Axin2 in acini obtained as in **B**. Number of mice=4. Student’s unpaired t-test. **G**. Same mammary gland tissues used in **Fig. 4B** were stained with CK14 antibody and DAPI. Graph shows MFI per acini. Number of mice=3/group. Graph represents mean± SEM. Unpaired student t-test. Scale bar=10μm. **H. *Nr2f1* KD does not affect proliferation in ECCs.** MMTV-HER2 early cancer cells were cultured in acini and transfected with either siControl or siNR2F1. After 48 h, acini were fixed and stained using an anti-pH3 antibody. Graph shows percentage of pH3 positive cells per acinus. Representative result of 3 independent experiments. Acini analyzed siControl=58; iNR2F1=39. Student’s unpaired t-test.

**Movies 1&2**- Early cancer acini transfected with siControl were imaged for 4 hours every 10 min.

**Movies 3&4**- Early cancer acini transfected with siNR2F1 were imaged for 4 hours every 10 min.

**Movies 5**- shControl early cancer cells injected into the fat pad of nude mice were imaged as described in Methods section for half an hour every 2 minutes using multiphoton confocal scope, 25x 1.05NA (XLPL25XWMP2, Olympus) water immersion objective. The movie is a representative region of one mouse.

**Movies 6**- shNR2F1 early cancer cells injected into the fat pad of nude mice were imaged as described in Methods section for half an hour every 2 minutes using multiphoton confocal scope, 25x 1.05NA (XLPL25XWMP2, Olympus) water immersion objective. The movie is a representative region of one mouse.

## Notes

### Competing Interest Statement

The authors have declared no competing interest.

